# Quench me if you can: Alpha-2-macroglobulin trypsin complexes enable serum biomarker analysis by MALDI mass spectrometry

**DOI:** 10.1101/2020.11.23.394759

**Authors:** Aleksandr S. Taraskin, Konstantin K. Semenov, Aleksandr V. Protasov, Alexey A. Lozhkov, Alexandr A. Tyulin, Aram A. Shaldzhyan, Edward S. Ramsay, Olga A. Mirgorodskaya, Sergey A. Klotchenko, Yana A. Zabrodskaya

## Abstract

One of the main functions of alpha-2-macroglobulin (A2M) in human blood serum is the binding of all classes of protease. It is known that trypsin, after such interaction, possesses modified proteolytic activity. Trypsin first hydrolyzes two bonds in A2M’s ‘bait region’, and the peptide ^705^VGFYESDVMGR^715^ is released from A2M. In this work, specifics of the A2M-trypsin interaction were used to determine A2M concentration directly in human blood serum using MALDI mass-spectrometry. Following exogenous addition of trypsin to human blood serum *in vitro*, the concentration of the VGFYESDVMGR peptide was measured, using its isotopically-labeled analogue (^18^O), and A2M concentration was calculated. The optimized mass spectrometric approach was verified using a standard method for A2M concentration determination (ELISA) and the relevant statistical analysis methods. It was also shown that trypsin’s modified proteolytic activity in the presence of serum A2M can be used to analyze other serum proteins, including potential biomarkers of pathological processes. Thus, this work describes a promising approach to serum biomarker analysis that can be technically extended in several useful directions.

## 1 INTRODUCTION

Alpha-2-macroglobulin (A2M) is one of the major blood plasma glycoproteins with a wide range of regulatory and transport functions [1–3]. One of its main functions is to inhibit a whole range of proteases, including trypsin (used in this work). Other A2M activities or features include: transport of cytokines, growth factors, and hormones; inhibition of complement system enzyme cascades and the kallikrein-kinin system; involvement in blood coagulation and fibrinolysis; and development of inflammatory reactions; and exhibition of immunosuppressive properties [4]. Foremost, A2M can form complexes with all classes of proteases. The large tetrameric complex of A2M binds a protease molecule so that the reactive center of the latter remains substantially free [1,5,6]. Therefore, proteases captured by A2M may hydrolyze low molecular weight synthetic substrates, as well as labile proteins without stringent secondary or tertiary structure [4,7,8]. Most proteins, including major serum proteins such as serum albumin and fibrinogen, are resistant to digestion by the trypsin-A2M complex. However, protamine sulfate peptide is hydrolyzed by this complex [7]. Moreover, it is known that, when complexed with A2M, trypsin becomes resistant to other serum inhibitors [2,7,9]. In addition to A2M, another major protease inhibitor is present in blood, α1-antitrypsin (A1AT), which also binds trypsin, but with a complete loss of the latter’s activity. The affinity of A2M for trypsin is 3-4-fold greater than the corresponding A1AT affinity (10^−13^ vs 5×10^−9^ M). Thus, A2M binds trypsin first. After depletion of trypsin, binding of A1AT proceeds until A1AT is completely inhibited. [10–14].

It is known that serum A2M concentrations vary considerably in a number of diseases [15]. Therefore, measurement of A2M may be valuable in diagnostics as well. Routine immunological or enzymatic methods are used to determine A2M concentration [16,17]. The former requires the use of specific antibodies, while the latter measures the hydrolysis rates of specific substrates. Moreover, antibody-based methods do not distinguish between the A2M free form and its reaction products. It should also be noted that the ability of antibodies to bind the antigen may vary dramatically depending on the conformation of latter. The conformation of A2M can change significantly, such as during interaction with proteases [4,18,19]. Given the large number of variables involved, it not clear that every method, including those commonly used, measure A2M in consistent ways.

To determine active human A2M serum concentrations, we propose the use of mass spectrometry (MALDI-MS) with an isotopically labeled internal standard. Our approach is based on peculiarities of the A2M/trypsin interaction, which results in the appearance of a characteristic peptide, VGFYESDVMGR [1]. This peptide is easily detectable by MALDI-MS, and its concentrations is proportional to A2M in blood serum [20].

In quantitative proteomics, mass spectrometric approaches have gained widespread use relatively recently. The most common approach is quantitative analysis of proteins using high performance liquid chromatography-mass spectrometry. However, this approach has a number of significant limitations associated with: the impossibility of performing quantitative multiplex analysis of several components at once; and the need for preliminary fractionation of samples. [21]. The method cannot be considered highly efficient and, in terms of analysis time, is inferior to MALDI (Matrix Assisted Laser Desorption/Ionization) mass spectrometry, which was used in this work [8].

## 2 MATERIALS AND METHODS

### 2.1 Human blood sera

Sera from 20 patients were kindly provided by the Department of Molecular Biology of Viruses (Smorodintsev Research Institute of Influenza, St. Petersburg): healthy donors – 5; patients with pyoinflammatory soft tissue diseases accompanied by sepsis – 5; patients with influenza A (H1N1 or H3N2), confirmed by RT-PCR (AmpliSens, Russia) – 5; and patients with COVID-19, confirmed by RT-PCR (“AmpliTest SARS-CoV-2”, Russian Centre for Strategic Planning) – 5. The presence of inflammation was confirmed using ELISA for detection of IL-6 and IL-10; INF-λ was also measured (section 2.5).

This work was carried out in accordance with the Code of Ethics of the World Medical Association (Declaration of Helsinki). Informed consent was obtained for experimentation with human samples or subjects.

### 2.2 Preparation of the ^18^O standard (^18^O-SYN-pep)

In order to permit quantitative analysis of the characteristic VGFYESDVMGR peptide in samples, its synthetic analogue was commercially synthesized (Verta Limited, St. Petersburg, Russia).

The ^18^O-labeled standard was prepared by isotopic exchange of ^16^O by ^18^O (located in carboxyl groups), according to our previously developed protocol [8,22]. One milligram of peptide was dissolved in 20 ul of 10% trifluoroacetic acid (TFA, Sigma-Aldrich), in an H_2_^18^O base (95-98% ^18^O, Cambridge Isotope Laboratories Andover, MA, USA), and incubated in a solid-state thermal block at 70°C, for 90 min. It was then dried in a SpeedVac vacuum evaporator (Eppendorf). The contents of the tube were dissolved in water and used as a standard for measurement of serum-derived peptide (**SER-pep**) concentrations (see 2.4). The precise concentration of the ^18^O-SYN-pep was determined using its absorbance at 280 nm (molar extinction coefficient ε = 1490 M^-1^·cm^-1^).

### 2.3 Direct tryptic hydrolyses of human blood serum for A2M assessment

Human blood serum, after dilution 25-fold, was mixed with equal volumes of modified, porcine trypsin (0.05 mg/ml stock in 50 mM bicarbonate buffer). After hydrolysis for several time periods at 20°C, reactions were stopped by adding equal volumes of 2% TFA. The concentration of the resulting peptide was assessed by isotopic ratio analysis via mass spectrometry.

### 2.4 Assessment of A2M concentration using MALDI mass spectrometry and ^18^O-SYN-pep

The trypsin-treated samples were mixed in equal volumes with ^18^O-SYN-pep of known concentration. The resulting mixture was mixed with HCCA matrix (Bruker) in equal volumes and spotted onto a target plate.

Mass spectra were recorded using the UltrafleXtreme MALDI-TOF/TOF instrument (Bruker Daltonics, Germany). The monoisotopic mass peak accuracy was 20 ppm. For quantitative measurements, each spectrum was a sum of 5,000 laser pulses. At least five spectra were recorded for each sample.

The concentration of SER-pep (VGFYESDVMGR peptide) was calculated, by formula **(1)**, from the ratio of peak areas corresponding to: ions associated with sample; and ions associated with ^18^O-SYN-pep (Fig. 1a). This is possible due to the fact that the isotope signals of the parent peptide and the standard do not overlap. Sample A2M concentrations were calculated from the known molar ratio: 1 mole A2M produce 2 moles of peptide.

**Figure 1.**
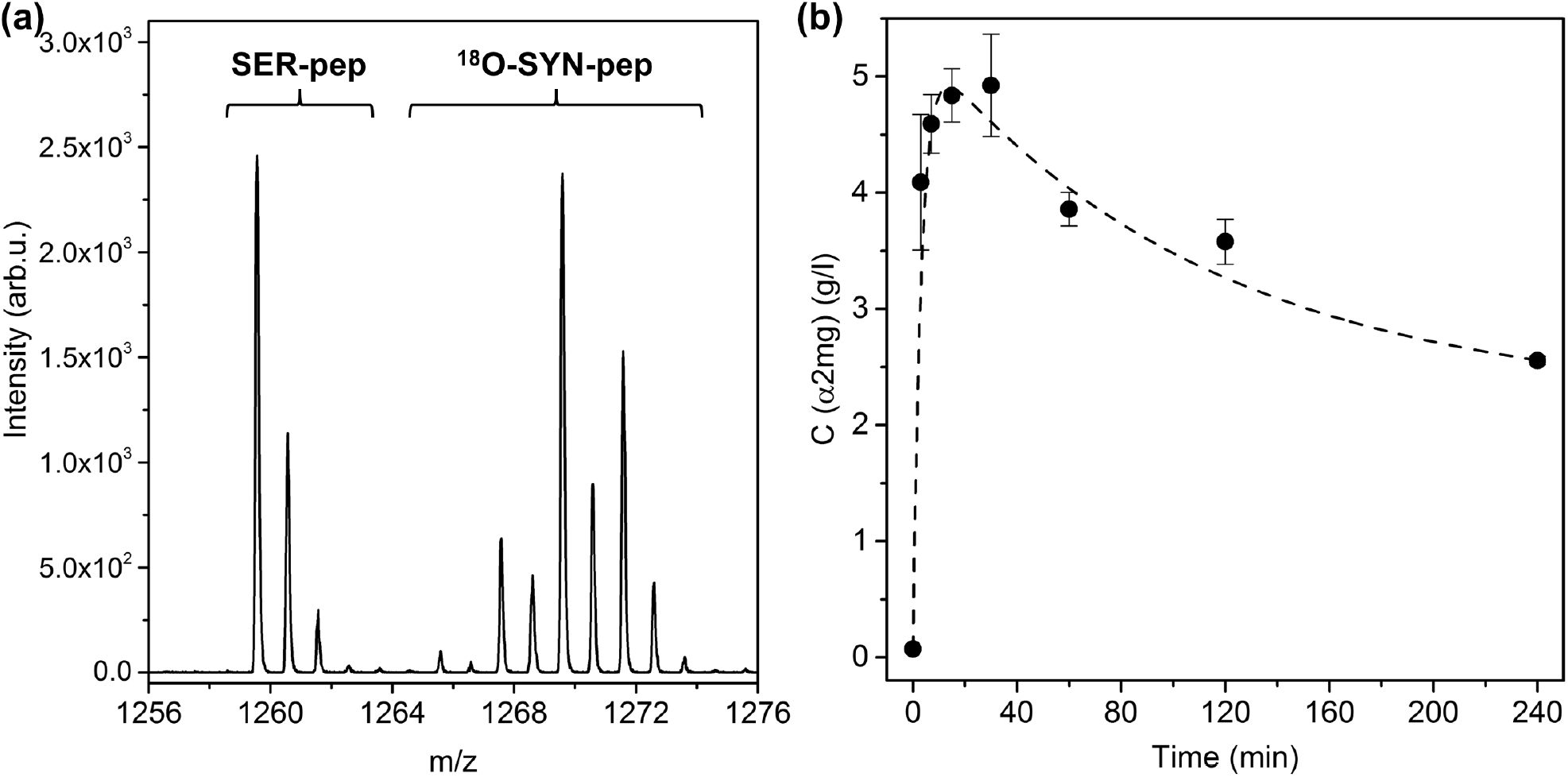
**(a)** MALDI mass spectrum of mixture containing the VGFYESDVMGR peptide (SER-pep) and the isotopically-labeled standard (_18_O-SYN-pep). **(b)** A2M concentration in healthy donor blood serum, depending on incubation time with added trypsin.

### 2.5 ELISA

ELISA, to determine A2M concentration in human blood serum, was performed according to manufacturer’s instructions (Ref K6610A, Immundiagnostik AG, Germany). Serum samples were diluted 1:50,000 with dilution buffer, in three steps, as recommended by the manufacturer. Absorption was measured with the CLARIOstar® microplate reader (BMG LABTECH, Germany) at 450 nm, with a 620 nm reference. Spline algorithm (linear regression, least square method) was used to calculate the results. The values obtained were multiplied by the dilution factor (50,000) to calculate actual concentration.

ELISAs for determination of interferon-λ (IFN-λ), interleukin-6 (IL-6), or interleukin-10 (IL-10) concentration were carried out using the following commercial kits or antibodies according to the manufacturer’s instructions. Lambda interferon was evaluated by Human IL-29/IL-28B (IFN-lambda 1/3) DuoSet ELISA (DY1598B, R&D Systems, USA). Interleukin-6 was measured by Human IL-6 DuoSet ELISA (DY206, R&D Systems, USA). Reagents for interleukin-10 analysis were: Rat Anti-Human IL-10 capture antibodies (554705, BD Biosciences, USA); Biotin Anti-Human and Viral IL-10 detection antibodies (554499, BD Biosciences, USA); and Recombinant Human IL-10 (1064-IL-010, R&D Systems, USA). In all cases, Costar EIA/RIA 8 well strips (cat. 2580, Corning, USA) were used to perform the analysis.

### 2.6 Statistical analysis

Data for A2M concentration comparison, obtained by two measurement methods (mass-spectrometry, *C*_*MS*_; ELISA, *C*_*ELISA*_), were processed using relevant statistical methods. A list of procedures performed is shown below.

1. The *C*_*MS*_ and *C*_*ELISA*_ sample values obtained were tested for homogeneity for each group (p<0.05) and outliers were removed (1 value from each sample of sizes 20 and 22 values) [23–25].
2. *C*_*MS*_(*C*_*ELISA*_*) dependence* was approximated using the least squares technique applied to a set of polynomials with ascending degrees beginning from zero. The lowest suitable degree was chosen using Fisher sequential goodness-of-fit test (p<0.05) [26,27] and developed related technique for non-random errors. As a result, an approximation in the form *C*_*MS*_ = *A* · *C*_*ELISA*_ + *B*, where A and B are fitted regression coefficients, was obtained.
3. To check the degree of linearity, one-sided confidence bounds for Pearson correlation coefficient [28,29], between *C*_*MS*_ and *C*_*ELISA*_ sample values, were calculated; these formed the interval [0.86; 1.00] (P=0.95). It is thus seen that strong correlation, in terms of the Chaddock scale, is present [30].
4. The confidence bounds were derived for coefficients A and B [31,32] to test their significance and to simplify the approximation curve, if permissible, as well as the developed technique for case of non-random and mixed-nature errors. The final estimate of *C*_*MS*_(*C*_*ELISA*_*) dependence* was obtained in the form *C*_*MS*_ = 2 · *C*_*ELISA*_. The Wald-Wolfowitz test [33,34] on the randomness of regression residuals was applied (p<0.05).
5. Using the regression *C*_*MS*_ = 2 · *C*_*ELISA*_, samples were normalized for representation on one unified scale (without MS/ELISA difference). The equality of their (*C*_*MS*_ and *C*_*ELISA*_) *means* was tested using the Student t-test [35–37] for two related samples. Equality of their *medians* (*P*((2 · *C*_*ELISA*_) < *C*_*MS*_) = 0.5) was tested using the Wilcoxon test with Iman’s correction [38,39] for related samples. These tests were successfully passed (p<0.05).
6. Finally, the standard deviation of random error (of the described mass-spectrometric method) was estimated for error related to conditions (different researcher or day, for example). The SD was 0.588 mg/ml; one-sided confidence bounds were [0, 0.878] mg/ml (P=0.95). We also estimated the ratio {error between and error MS} with the MALDI-MS instrumental error with the obtained total random error: it causes not less than 52% in the mentioned measurement random variability.

### 2.7 Isolation of alpha-2-macroglobulin

Gel filtration. The sera from two healthy donors (1.5 ml) were diluted two-fold and concentrated (to 1 ml final volume) using Vivaspin Turbo 15 (Sartorius) units. To isolate A2M, the AKTA pure L chromatographic system (GE Healthcare) was used. The column, HiLoad 26/60 Superdex 200 pg (GE Healthcare), was equilibrated with 240 ml of PBST (Sigma) at a flow rate of 30 cm/h. The concentrated serum was introduced into the column through a 1 ml injection loop. Elution was performed using the same buffer, with optical density monitoring at 280 nm (OD_280_). Fractions (2.5 ml) were collected in automatic mode upon A_280_ values above 10 mAU.

The same protocol was used for standard proteins with known molecular weight but without fraction collection. Fractions collected (retention volume *V*_8_ = 100 ÷ 150*ml*) were analyzed using SDS-PAGE, as described [40]; staining was performed with colloidal Coomassie [41]. Fractions containing A2M tetramers were combined and concentrated to 1 ml final volume (Vivaspin Turbo 15, Sartorius).

Metal-affinity chromatography. A HisTrap FF chromatographic column (GE Healthcare), with a volume of 1 ml and Zn(II) charged, was used. It was equilibrated with 5 ml of start buffer (50 mM sodium phosphate, 300 mM sodium chloride, 1 mM imidazole). The A2M sample, obtained earlier, was introduced into the column through a 1 ml injection loop. Next, the column was washed with 5 ml of start buffer, and the protein was eluted with 5 ml of elution buffer (50 mM sodium phosphate, 300 mM sodium chloride, 200 mM imidazole). The protein was collected and transferred into PBS using a Bio-Scale Mini P6 chromatographic cartridge (Bio-Rad) with a volume of 10 ml. The protein concentration (*C*, g/l) was determined, at the desalting stage, from the peak area (*A*_*280*_), as follows: *C* = *M*_*r*_*A*_280_/*εl*, where monomer molecular weight *M*_*r*_ was taken to be 181 kDa; optical path length *l* = 1*cm*; the A2M monomer’s theoretical molar extinction coefficient *ε* =145440*M*^−1^*cm*^−1^ was calculated using human A2M amino acid sequence P01023 [42] at a Prot-Param tool [43], assuming all pairs of Cys residues form cysteines. Sample purity was monitored by SDS-PAGE [40] followed by densitometry of the colored bands using ChemiDoc XRS software (BioRad).

### 2.8 Protein identification using MALDI MS

Stained protein bands were cut out, and tryptic hydrolyses were performed as described [44]. Mass spectra of tryptic peptides were registered as described in section 2.4. Protein identification was performed using MASCOT (http://www.matrix-science.com) and the Swiss Prot database (https://www.uniprot.org). The oxidation of methionine was marked as a variable modification. Monoisotopic mass measurement accuracy was limited to 20 ppm. Identification of individual peptides (observed in mass spectra) was performed in tandem configuration mode (msms). In that mode, the monoisotopic mass measurement accuracy (of the fragments) was 0.2 Da. Protein identification were performed as described above.

## 3 RESULTS AND DISCUSSION

### 3.1 Basic principles

In our previous study [20], the possibility of A2M concentration assessment, directly in human blood serum, was demonstrated. Briefly, it was shown that: after addition of trypsin to serum and a few minutes of incubation, an intense, singly-protonated quasimolecular ion (*m*/*z*=*1259*.*57*) is registered in the MALDI mass spectrum. Detailed analysis in tandem ion registration mode (msms) showed that the observed ion corresponds to peptide from human alpha-2-macroglobulin (Score 59/42). Taking in account the peculiarities of the A2M-trypsin interaction [1], the discovered peptide (^705^VGFYESDVMGR^715^, ***SER-pep***) was found to be a fragment of the so-called ‘bait region’ of A2M (a.a.r. 690-728), as expected; this region is involved in trypsin binding and inactivation by A2M in blood serum. It was also shown that the amount of peptide detected is equal to the amount of trypsin bound with A2M [20].

It should be noted that, depending on the amount added to blood serum, trypsin can exist in one of three possible states: in a complex with A2M; in a complex with A1AT; or in a free form (when its concentration exceeds the binding capacity of A2M and A1AT). In order to determine A2M concentration, all A2M molecules should be occupied by added trypsin (trypsin reagent added exogenously to serum). Any trypsin exceeding the binding capacity of A2M is completely inhibited by A1AT and, as such, does not influence production of the characteristic peptide (VGFYESDVMGR peptide). It is known that each A2M tetramer binds two trypsin molecules [10], thus it should also produce two characteristic peptide molecules. It was assumed that *C*_*A2M*_ = *C*_*SER-pep*_/2. The aforementioned principles are the basis for the methods developed in this work.

### 3.2 Kinetics of SER-pep release upon trypsin exposure

To develop a precise and reliable method, determination of the ideal trypsin/serum incubation time was necessary. For this purpose, a kinetic experiment was performed (Fig. 1b). Following the previously-described experimental method [20], 25-fold diluted serum was incubated with 0.05 mg/ml trypsin (a 10-fold molar excess which binds all available bait regions). Reactions were stopped after 0 (control), 3, 7, 15, 30, 60, 120, or 240 minutes. Concentrations of SER-pep were then measured, as described in the Materials and Methods, using internal standard (synthetic VGFYESDVMGR peptide) labeled with ^18^O (^***18***^***O-SYN-pep***).

It has been shown that the effectiveness of MALDI ionization, for the peptide and its isotopically-labeled standard, are the same [22]. Thus, the total intensity of the isotope distribution peak (area) of SER-pep 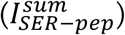, or of the ^18^O-SYN-pep 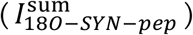, can be considered proportional to their molar concentrations (*C*_*SER-pep*_and *C*_180-*SYN-pep*_ respectively):

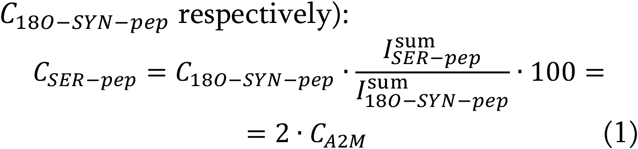

It should be noted that the mass spectrum ion intensity of ^*18*^*O-SYN-pep* and serum-derived pep tide must be similar for good analysis accuracy.

According to the *SER-pep* concentration dynamics (Fig. 1b), it can be concluded that the highest serum SER-pep concentration is observed after 10-30 minutes of incubation with trypsin. After 30 minutes, a decrease in peptide concentration is seen; this may be due to interaction with other serum components. Based on this data, 15 minutes was selected as the optimal incubation time for the analysis method. It should be noted that the selected time was valid for a wide range of A2M concentrations, and there was no significant difference between performing the experiment at room temperature or at 37°C.

### 3.3 Verification of the developed method using ELISA

Commercial ELISA reagents were used to verify the method developed (for determination of blood serum A2M concentration using mass spectrometry and ^18^O-SYN-pep). For this purpose, 20 blood serum samples were used: 5 from healthy donors; 5 from patients with sepsis; 5 from patients with influenza A; and 5 from patients with COVID-19. This sample size meets the minimum requirements for statistical analysis. In each case, A2M concentration was measured by two methods: MALDI MS with ^18^O-SYN-pep; and ELISA with A2M antibodies.

It is generally accepted to classify A2M as an acute phase protein whose production increases in response to secretion of the proinflammatory cytokine IL-6 [45]. Thus, concentrations of proinflammatory cytokines (IL-10, IL-6), along with IFN-λ, were also measured to provide unequivocal laboratory evidence of inflammation (Table 1). We observed an increase in IL-6 level, which can also be elevated with sepsis, influenza, and SARS-CoV-2 [46–48]. The level of IL-10 was also increased here, the excess production of which is associated with the development of immunopathology [49]. Elevated IL-10 has been observed with SARS-CoV-2 [48]. The greatest differences, among our groups, were observed for the sepsis group. Lambda interferon is capable of stimulating IL-10-dependent IL-6 secretion by PBMCs [50]. Elevated IFN-λ was not detected, even in the sepsis group. This further emphasizes the insignificant contribution of IFN-λ to development of “cytokine storm” or generalized acute inflammatory processes [51].

**Table 1.** Comparative biomarker analysis. Serum A2M concentrations are shown, as determined using mass spectrometry or ELISA. ELISA-determined values are also shown for IFN-λ, IL-10, and IL-6. The groups are: (1) healthy donors (“Control”); (2) patients with pyoinflammatory soft tissue disease accompanied by sepsis (“Sepsis”); (3) those with influenza A(H1N1) or (A)H3N2 (“Influenza A”); and (4) those with COVID-19.

| Patient | MS | ELISA |  |  |  |
| --- | --- | --- | --- | --- | --- |
| | A2M (g/l) | A2M (g/l) | IFN-λ (ng/l) | IL-10 (ng/l) | IL-6 (ng/l) |
| Control |  |  |  |  |  |
| 1 | 1.53 | 0.73 | 0.0 | 1.3 | 0.0 |
| 2 | 2.83 | 1.66 | 28.5 | 3.1 | 2.8 |
| 3 | 1.71 | 0.95 | 0.0 | 0.0 | 0.0 |
| 4 | 4.46 | 2.42 | 0.4 | 0.0 | 0.0 |
| 5 | 5.50 | 2.09 | 0.0 | 0.0 | 0.0 |
| Sepsis |  |  |  |  |  |
| 6 | 6.42 | 2.67 | 0.0 | 85.9 | 571.7 |
| 7 | 8.56 | 2.45 | 0.0 | 41.4 | 3.2 |
| 8 | 1.72 | 1.23 | 0.0 | 47.3 | 38.4 |
| 9 | 1.05 | 0.81 | 0.0 | 52.3 | 1.3 |
| 10 | 3.44 | 1.30 | 0.0 | 103.8 | 3343.9 |
| Influenza A |  |  |  |  |  |
| 11 | 3.99 | 2.19 | 0.0 | 9.5 | 0.5 |
| 12 | 2.01 | 1.37 | 0.0 | 13.8 | 20.5 |
| 13 | 7.18 | 3.29 | 5.2 | 9.1 | 64.2 |
| 14 | 5.58 | 3.02 | 0.0 | 6.1 | 3.5 |
| 15 | 1.72 | 0.69 | 0.0 | 12.6 | 9.4 |
| COVID-19 |  |  |  |  |  |
| 16 | 2.76 | 1.60 | 0.0 | 1.7 | 0.0 |
| 17 | 4.15 | 2.33 | 0.0 | 7.1 | 2.8 |
| 18 | 1.81 | 1.03 | 0.0 | 7.3 | 14.6 |
| 19 | 2.44 | 1.27 | 0.0 | 6.5 | 3.5 |
| 20 | 1.83 | 1.01 | 0.0 | 20.0 | 24.2 |

Normally, A2M levels in human blood serum vary from 1 to 6 g/l, depending on age and sex [4,52]. In the data presented (Table 1), normal A2M levels were generally observed with both methods (mass spectrometry and ELISA), despite elevated IL-10 and IL-6 in some cases. This indicates the presence of inflammatory processes that are not correlated with elevated A2M concentration; this feature has been noted by other authors [53]. It should be emphasized that the aim of this experiment was to explore if there is a difference in A2M concentration, as determined by MS (*C*_*MS*_) or ELISA (*C*_*ELISA*_). At first glance, it may seem that *C*_*ELISA*_ is lower in all cases than *C*_*MS*_. To formally analyze such a conclusion, statistical analysis was performed wherein the two data types (A2M concentration in g/l, determined by *C*_*MS*_ or *C*_*ELISA*_)) were compared. A scatter plot (*C*_*MS*_ (*C*_*ELISA*_) dependence) and a regression curve (of the form *C*_*MS*_ =*A* · *C*_*ELISA*_ + *B*, where *A* =2.142 ± 0.430, *B* =−0.1745 ± 0.7793; 95% confidence intervals) were produced (Figure 2).

**Figure 2.**
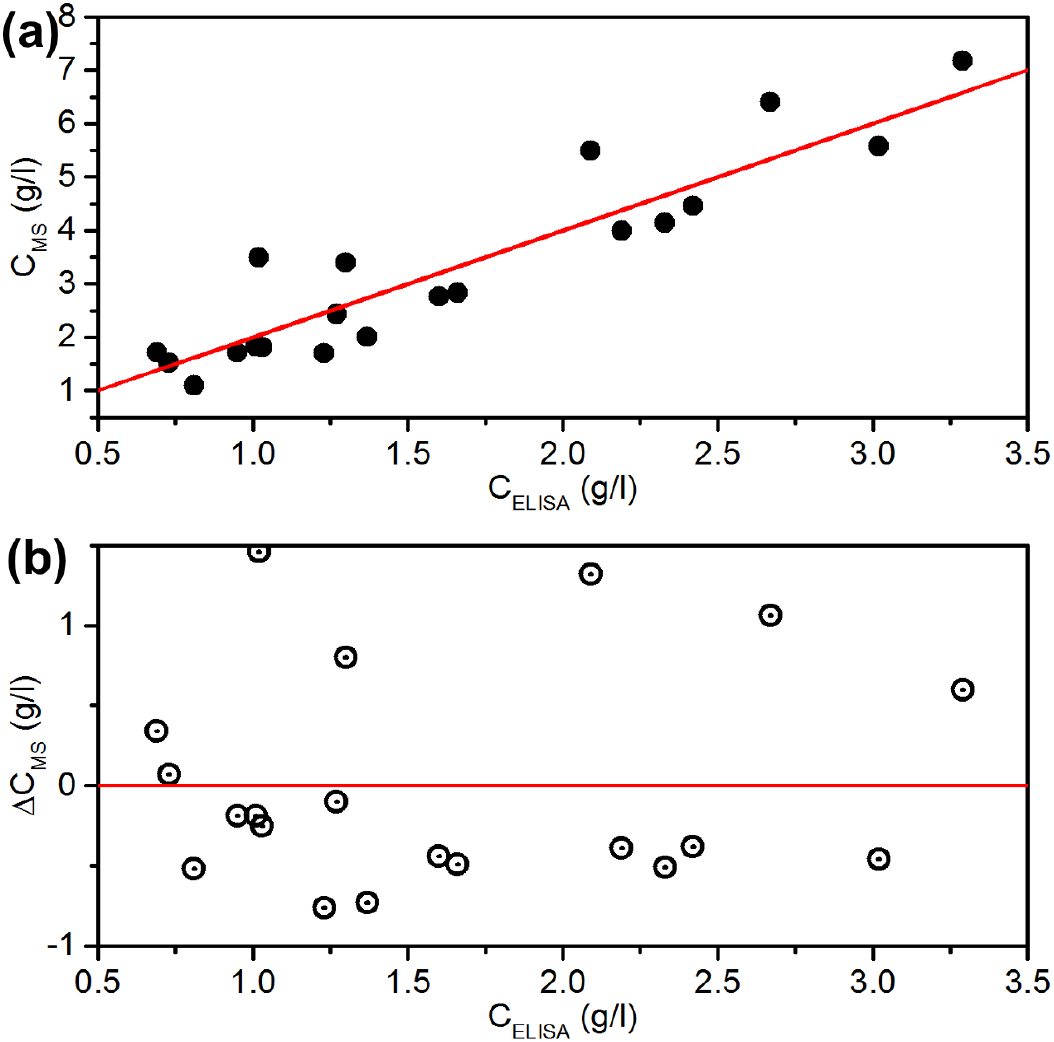
**(a)** Scatter plot (black circles) and regression curve (red line) C_^__ = 2 · C_‘ab_c_. **(b)** Residuals.

Taking into account the applied statistical criteria, several arguments can be made. Firstly, there is no reason to believe that the value of the coefficient *A* differs from two, or the coefficient *B* – from zero, i.e. *C*_*MS*_ =2 · *C*_*ELISA*_. Secondly, there is no reason to believe that the regression residuals (the differences between the points (*C*_*MS*_, *C*_*ELISA*_) and the values of the regression curve *C*_*MS*_(*C*_*ELISA*_)) behave in a non-random way. The correlation coefficient between the values, with a probability of 0.95, lies in the [0.83; 1.00] interval (one-sided confidence interval with right limit fixed at a value of +1). Thus, it was determined that there is an unambiguous correlation between A2M concentrations determined by MS and by ELISA. However, the ELISA reagents used yield A2M concentrations exactly two times lower than MALDI MS with ^18^O-SYN-pep.

It should be noted that if trypsin is added to serum before ELISA analysis (at the same trypsin concentration used for MS analysis), *C*_*ELISA*_ values become higher and closer to *C*_*MS*_ values (Table 2). This fact indicates that low *C*_*ELISA*_ may be related to structural features of A2M antigen/antibody interactions at the core of the ELISA analysis mechanism. It is known that A2M undergoes significant conformational changes during trypsin binding [4,18,19]. Such changes may lead to modification of some epitopes, with the latter becoming more antibody accessible. That would explain changes in antigen/antibody interaction and, as a consequence, the increase in *C*_*ELISA*_. Moreover, it has been established that A2M structure is acutely sensitive to freezing and lyophilization, which is a crucial factor with lyophilized A2M standards [54].

**Table 2.** Influence of serum trypsin-pretreatment on A2M concentrations determined by ELISA. Values determined by MS are normalized using the ELISA serum dilution factor.

| № | Alpha-2-macroglobulin concentration (ng/ml) |  |  |
| --- | --- | --- | --- |
|  | ELISA without trypsin | ELISA with trypsin | Mass Spectrometry |
| 1 | 58.7 | 80.2 | 85.4 |
| 2 | 109.3 | 130.9 | 159.9 |
| 3 | 84.8 | 108.7 | 117.6 |

This is relevant because the commercial ELISA kit used here (Ref K6610A, Immundiagnostik AG, Germany), according to manufacturer’s instructions, features an A2M standard that, after reconstitution, is stored at -20°C before use in analysis. This may also influence A2M/antibody interaction capability.

To verify that the developed method measures true A2M concentration, pure A2M was isolated from human blood serum by: gel-filtration chromatography (Fig. S1); metal-affinity chromatography with Zn(II) (Fig. S2,a); and desalting (Fig. S2,b); followed by storage at +4°C. The final A2M concentration was calculated as described in Material and Methods section 2.7. The identity of the purified protein with human A2M was confirmed by MALDI MS (Score/threshold is 368/66) after PAGE (Fig. 3) and trypsin digestion.

**Figure 3.**
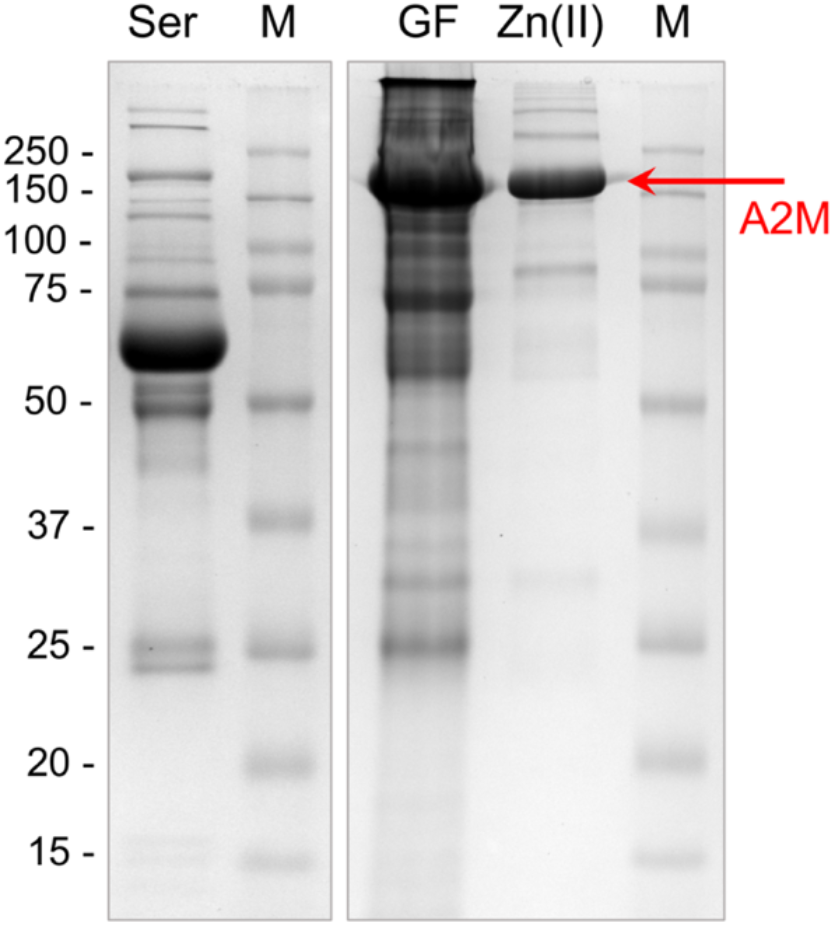
**All stages of A2M chromatographic isolation viewed by PAGE**. Ser – the initial human blood <u>serum</u>, diluted 40-fold; GF – combined fractions after <u>gelfiltration</u>; and Zn(II) – combined fractions after metal-affinity chromatography with <u>Zn(II)</u>, containing pure A2M (marked by arrow), which was reliably-identified using MALDI mass spectrometry. M – molecular weight marker.

The ‘true’ A2M concentration, accordingly to chromatography, was **0. 8** g/l. Using MALDI MS with ^18^O-SYN-pep, the concentration determined was **0. 80** ± **0. 06** g/l. In parallel, ELISA indicated an A2M concentration of **0. 42** ± **0. 02** g/l. Thus, the same tendency, noted earlier with human blood serum, is seen. This fact also confirmed that the developed method determines true A2M concentration.

### 3.4 Potential application for other serum biomarkers

As mentioned above, following A2M/trypsin interaction, trypsin doesn’t completely lose activity. Rather, it is modified: trypsin retains the proteolytic ability to cleave low-molecular-weight polypeptides (< 20 kDa) [55]. Since the proteolytic activity of trypsin in complex with A2M is limited, and it loses the ability to hydrolyze major serum proteins (for example, albumin) due to their large sizes (well above 20 kDa), such A2M-trypsin complexes may be useful as a research tool for identifying minor, small serum proteins. It has been shown earlier that serum amyloid A (SAA) fragments can be detected in mass spectra after serum treatment with trypsin for two hours [8,20]. Thus, it was observed in our experiments that some other ions, that differ from SAA, appear in the mass spectrum with longer incubation of serum with trypsin (marked by asterisk in Fig. 4).

**Figure 4.**
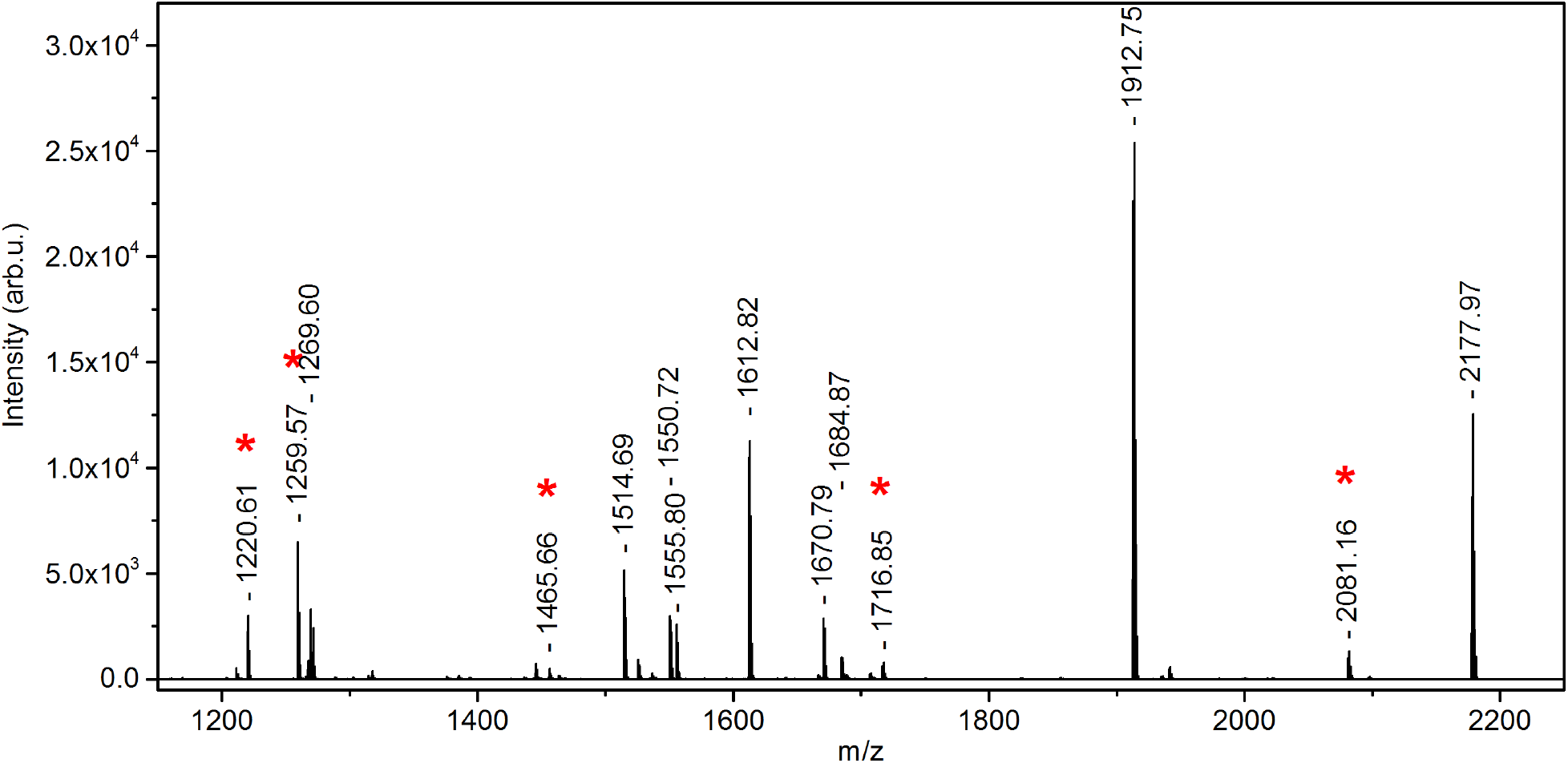
MALDI mass spectrum of serum treated with trypsin for two hours. Some identified ions, which differ from SAA, are marked (*).

The observed peptides (marked by asterisk) were reliable identified using tandem mass spectrometry (Table 3). Thus, it was shown that trypsin in complex with A2M produces peptides detectable by MALDI MS.

**Table 3.**
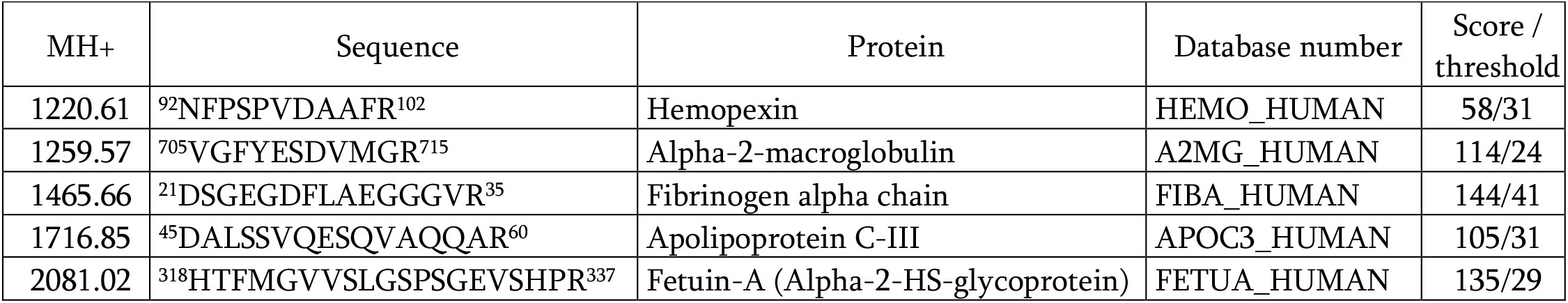
Identified peptides.

This finding makes it possible to consider the trypsin-A2M complex, artificially formed in blood serum, as a tool for determining the concentrations of various serum proteins without preliminary fractionation. Adaptation of the approach described here (for A2M) to enable detection of Examples include: fetuin-A concentrations correlate with disease severity in sepsis patient [56,57]; and the levels of some SAA variants (SAA1, SAA2) in human blood serum are dramatically elevated (up to 10^5^-fold) in patients with acute phase inflammation [58]. It seems plausible that the spectrum of potential biomarkers may turn out to be much wider if other ranges of peptides, induced (by A2M-trypsin-complex proteolytic activity) in the sera of patients with other diseases (not considered here), are analyzed.

## 4 CONCLUSIONS

In this work, a new method for determining alpha-2-macroglobulin concentration, directly in blood serum using an isotopically-labeled standard and MALDI mass spectrometry, was optimized and verified. This technique paves the way for quantification of proteins, including potential markers of various pathological conditions, without preliminary fractionation of blood serum. This is enabled by the modified proteolytic activity profile of trypsin when in the presence of A2M.

We acknowledge that different diagnostic methods and reagents all have specific features that promote or hinder reliable, precise measurement. Here, we noted lower values with the selected ELISA reagents. Upon treating serum in an analogous manner prior to ELISA, the difference became much smaller. The correlation between ELISA-based and MS-based values was consistent and supported by statistical analysis. Other commercial reagents may yield different values or trends. A notable benefit of the approach here is that it depends on stable, synthetic peptide, not labile reagents. Freezing, thawing, lyophilization, etc., are important factors that affect all commercial standards (typically for ELISA). Handling conditions for the synthetic peptide, used here for mass spectrometry, are unlikely to provoke large errors in MS-derived data, as the fundamental mechanism is not based on antibody-antigen interaction or associated structural influences in that system.

The techniques described here can likely be extended to permit analysis of many proteins that yield fragments detectable by mass spectrometry. The range of potential biomarkers that may be quantifiable (by means of A2M-trypsin complex interaction), directly in blood serum, is quite wide. It is especially promising that such approaches can simultaneously measure selected biomarkers.

## AUTHOR CONTRIBUTIONS

*A.S.T*. – Validation, Investigation, Writing - Original Draft; *K.S*. – Formal analysis, Writing - Original Draft; *A.P*. – Investigation; *A.L*. and *A.S*. – Investigation, Writing - Original Draft; *A.A.T*. – Validation, Investigation; *E.R*. – Writing - Original Draft, Writing - Review & Editing; *O.M*. – Conceptualization; *S.K*. – Resources; *Y.Z*. – Investigation, Writing - Original Draft, Visualization.

## ACKNOWLEDGMENTS

This work was supported by a Russian state assignment for fundamental research, No 0784-2020-0023 (IL-6, IL-10, INF-λ assays and analyses); by a Russian Science Foundation grant No 19-71-00127 and by the Ministry of Science and Higher Education of the Russian Federation as part of World-class Research Center

## SUPPLEMENTARY MATERIALS

**Figure S1.**
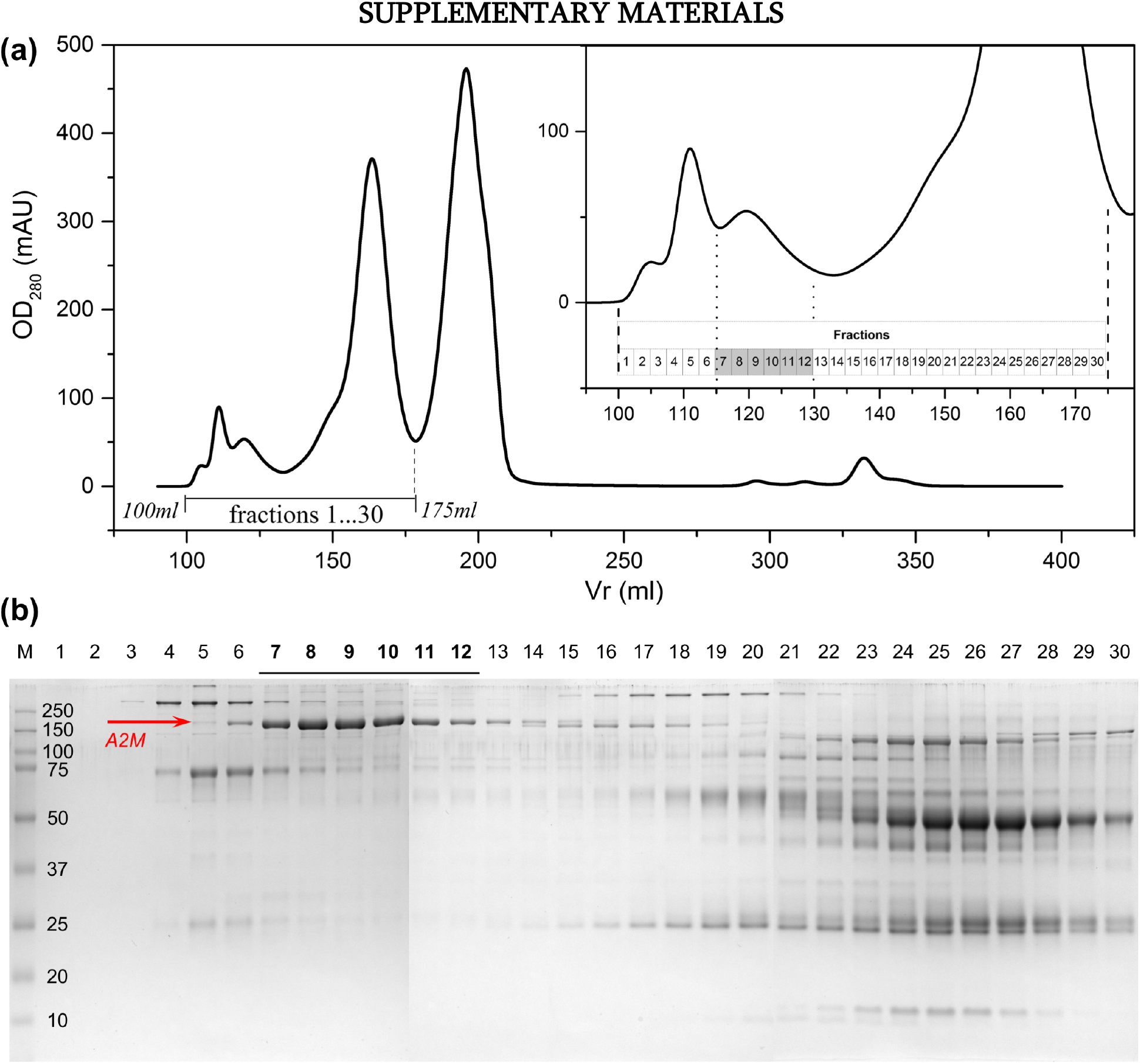
First step A2M isolation from human blood serum. **(a)** Gel filtration, with optical density at 280 nm (OD_280_) vs. retention volume (Vr). With the calibrated protein set used, A2M tetramers were expected to appear in the 110 - 140 ml retention volume range. All fractions collected were analyzed using PAGE **(b)**. A2M-containing fractions (numbers 7-12, marked by gray highlights and arrow) were combined.

**Figure S2.**
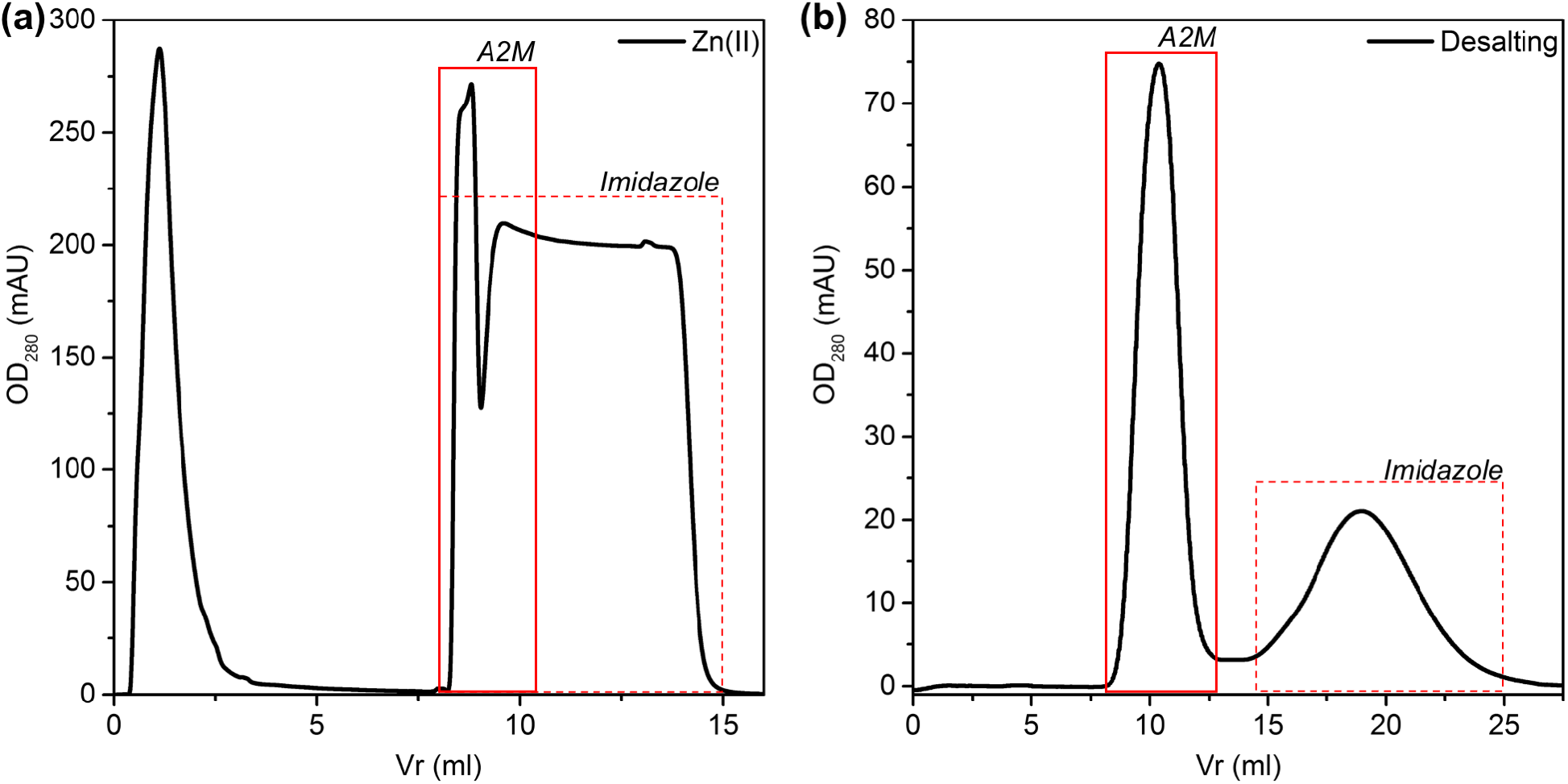
Second step A2M isolation from human blood serum: **(a)** metal affinity chromatography, followed by **(b)** the desalting stage. Fractions of interest (marked with red square) were combined.

